# Seizures cause sustained microvascular constriction associated with astrocytic and vascular smooth muscle Ca^2+^ recruitment

**DOI:** 10.1101/644039

**Authors:** Cam Ha T. Tran, Antis G. George, G. Campbell Teskey, Grant R. Gordon

**Author notes:** Corresponding Author: Dr. G. Campbell Teskey, Department of Cell Biology and Anatomy, Cumming School of Medicine; University of Calgary, 3330 Hospital Drive N.W., Calgary, Alberta, Canada, T2N 4N1. These authors contributed equally to this work (G. Campbell Teskey, Grant R. Gordon).

## Abstract

Previously we showed that seizures result in a severe hypoperfusion/hypoxic attack that results in postictal memory and behavioral impairments (Farrell et al., 2016). However, neither postictal changes in microvasculature nor Ca^2+^ changes in key cell-types controlling blood perfusion have been visualized *in vivo*, leaving essential components of the underlying cellular mechanisms unclear. Here we use two-photon microvascular and Ca^2+^ imaging in awake mice to show that seizures result in a robust vasoconstriction of cortical penetrating arterioles, which temporally mirrors the prolonged postictal hypoxia. The vascular effect was dependent on cyclooxygenase-2, as pre-treatment with ibuprofen prevented postictal vasoconstriction. Seizures caused a rapid elevation in astrocyte endfoot Ca^2+^ that was confined to the seizure period. Vascular smooth muscle cells displayed a significant increase in Ca^2+^ both during and following seizures, lasting up to 75 minutes. The temporal activities of two cell-types within the neurovascular unit lead to seizure-induced hypoxia.

**Highlights:**

- Seizures lead to equivalent levels of postictal hypoxia in both male and female mice
- Calcium elevation in astrocyte endfeet is confined to the seizure
- Postictal vasoconstriction in awake mice is mediated by cyclooxygenase-2
- Calcium elevation in vascular smooth muscle cells is enduring and correlates with vasoconstriction.

## Introduction

*Farrell et al., (2016)* systematically investigated local oxygen levels and blood flow following evoked and self-generated seizures in behaving rodents and discovered a severe hypoxic event (pO_2_ < 10 mmHg) that lasted over an hour. This phenomenon was the result of blood hypoperfusion and it generalized to people with epilepsy (Farrell et al., 2016; Gaxiola-Valdez et al., 2017; Li et al., 2019). Furthermore, *Farrell et al., (2016)* demonstrated that brain region-specific postictal memory and behavioral impairments were the consequence of this hypoperfusion/hypoxic event. Thus, seizures can result in a stroke-like attack which is responsible for the postictal state and may also be implicated in chronic behavioral comorbidities and anatomical alterations associated with epilepsy (Farrell et al., 2017).

Cyclooxygenase (COX)-2 and L-type Ca^2+^ channel activity during seizures were identified as key mechanisms necessary for the precipitous drop in tissue oxygen (Farrell et al., 2016). Using inhibitors of those targets it was demonstrated that the block of postictal hypoxia was downstream of epileptiform neural activity, pointing to direct actions on contractile components of the neurovascular unit. However, previously employed oxygen sensitive probes and laser Doppler flowmetry do not provide information on the activities of the cell-types underlying the phenomenon. For example, vascular smooth muscle cells (VSMCs) coat penetrating and pial arterioles (Grant et al., 2019; Hill et al., 2015). VSMCs are the end effectors controlling arteriole diameter and make a significant contribution to the regulation of micro-circulatory blood flow and cortical oxygenation (Nishimura et al., 2007; Sakadžić et al., 2016). VSMC membrane potential, coupling to L-type Ca^2+^ entry for excitation-contraction coupling, is a central mechanism mediating changes to arteriole tone. However, Ca^2+^ independent mechanisms of vasoconstriction also exist (Neppl et al., 2009; Toda, 1982), even during pathological vasospasms (Sato et al., 2000); thus, it is important to measure VSMC Ca^2+^ activity during and after a seizure to corroborate or refute a vascular Ca^2+^ hypothesis. Previous amelioration of postictal hypoxia with nifedipine argues in favour for a contribution from sustained VSMC free Ca^2+^ (Farrell et al., 2016). Additionally, astrocytes are important players in the regulation of local brain blood flow via their peri-vascular endfeet, which communicate to VSMCs. Astrocytes can release a number of vasodilator or vasoconstrictor messengers in a Ca^2+^ dependent manner (Howarth, 2014; Mishra, 2017). While there is little doubt that a large, transient increase in endfoot Ca^2+^ can control arteriole diameter (Gordon et al., 2008; Mulligan & MacVicar, 2004; Srienc et al., 2012; Straub et al., 2006), recent demonstrations also showed that endfeet can undergo long-lasting changes to free Ca^2+^ in response to neural activity, which translated into enduring increases in steady-state arteriole tone (Mehina, Murphy-Royal, & Gordon, 2017). Additionally, quantitative elevations in astrocyte endfeet Ca^2+^ correlate with microvascular diameter changes during epileptiform activity using the 4-AP seizure model in anesthetized mice (Zhang et al., 2019). However, examining Ca^2+^ signals in astrocyte endfeet and VSMCs in awake mice, rather than in brain slices or *in vivo* preparations utilizing anesthesia or sedation, is important because signaling pathways involving neurons, astrocytes and microvascular can crosstalk in a realistic manner. Thus, imaging behaving mice is important to prevent misrepresentation of the relationships between hypersynchronous neural activity, astrocyte and VSMC Ca^2+^, and hemodynamics (Tran & Gordon, 2015b, 2015a; Tran, Peringod, & Gordon, 2018). Here we performed *in vivo* two-photon imaging on awake head restrained mice to examine the Ca^2+^ activity patterns of cortical astrocytes and VSMCs during ictal and postictal periods to gain insight into the cellular underpinnings of seizure induced hypoperfusion/hypoxia.

## Results

### Long-lasting hypoxia follows maximal electroconvulsive seizures in awake freely moving mice and generalizes across sexes and cre-lox lines

We previously demonstrated seizure induced hypoxia using several animal models of epilepsy (Farrell et al., 2016), but for the purposes of examining the phenomenon in head restrained active mice under the two-photon microscope, we employed Maximum Electroconvulsive Seizures or MES for this study (Young et al., 2006). This was because MES evokes time-locked and wide spread seizure activity across the neocortex (Bondy et al., 1987), which can be captured at any arbitrary penetrating arteriole. Before imaging, we first validated MES-induced hypoxia in male and female C57Bl/6 mice that were awake and freely moving. Local field potentials and the partial pressure of oxygen (pO_2_ in mmHg) were chronically monitored in layer V of mouse barrel neocortex over 3 brief and discrete seizures (Figure 1A). Two hundred milliseconds of MES reliably manifested hypersynchronous epileptiform activity (Figure 1A) lasting between 32 to 57 seconds for males and 33 to55 seconds for females (Figure 1B,C inset), which produced tonic-clonic behavioural seizures analogous to generalized bilateral seizures in people with epilepsy. The mean baseline partial pressure of oxygen in the barrel cortex of freely-moving mice over the 2 days of recordings ranged between 20 and 30 mmHg in males and 18 and 32 mmHg in females (p>0.05)(Figure 1B,C). After each seizure, oxygen levels fell rapidly, taking over an hour to recover to baseline levels. Following the third seizure, a sustained decrease in pO_2_ was observed that was below the severe hypoxic threshold of 10 mmHg (Figure 1B,C). This threshold is defined with a molecular, cellular and clinical signature (Erecińska & Silver, 2001; Höckel & Vaupel, 2001; Jiang et al., 2009; van den Brink et al., 2000) and we quantified the integral (area) of the pO_2_ curve below this oxygen level (Farrell et al., 2016). Since no significant differences in seizure duration and postictal hypoxia were observed between the two sexes (Figure 1D), we mixed both sexes for subsequent experiments and pooled the data. Next, we tested for similar MES-induced hypoxia in the knock-in cre-lox crosses planned for two-photon fluorescence microscopy. First, postictal hypoxia was examined in *Slc1a3*-Cre/ERT x RCL GCaMP3 mice, which permit visualization of astrocyte free Ca^2+^ using the Excitatory Amino Acid Transporter 2 promoter (or GLAST)(Slezak et al., 2007). Second, we examined *Pdgfrβ*-Cre x RCL-GCaMP6s mice, which permits visualization of mural cell free Ca^2+^ using the Platelet derived growth factor receptor beta promoter (Cuttler et al., 2011). The MES elicited similar postictal hypoxia measures (Figure 1E-G) across both cre-lox cross lines and compared to C57Bl/6J mice. These data show that MES induces reliable seizures and that the phenomenon of postictal hypoxia generalizes across sexes and different mouse lines.

**Figure 1:**
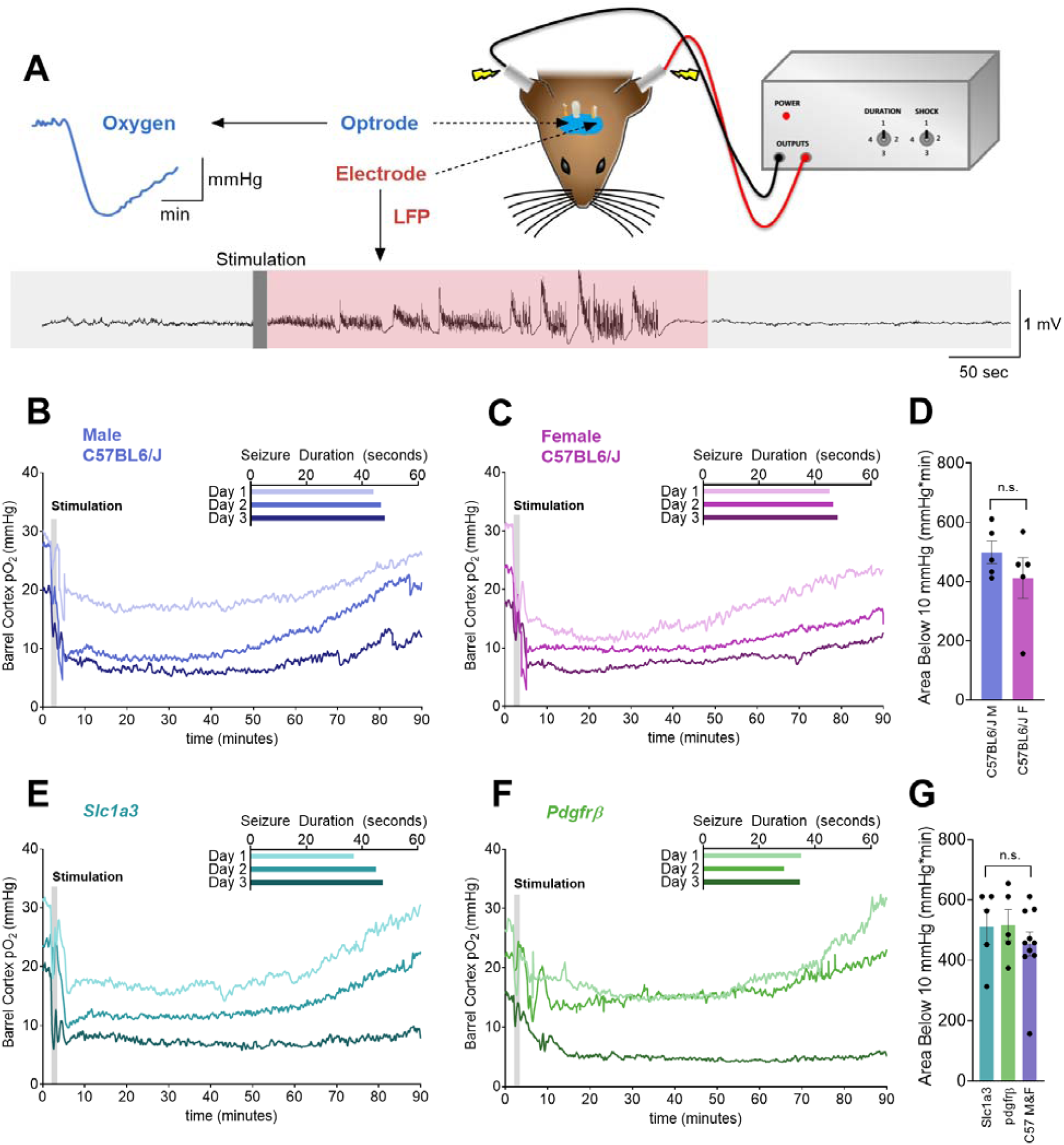
MES seizures induce postictal hypoxia that generalizes across sex and strains. (**A**) Experimental setup has mice with a chronically implanted optrode and electrode in their barrel cortex. Awake freely moving mice received one 0.2 sec MES per day for three days with concurrent local field potential and local partial pressure of oxygen (pO_2_) recordings. (**B and C**) Mean oxygen profile before, during, and after MES in male (*B*) and female (*C*) C57BL6/J mice (n=5, each). The inset shows the duration of electrographic seizures over the 3 seizures. (**D**) Quantification (mean ± SEM) of the area (depth and duration) below the severe hypoxic threshold (pO_2_<10mmHg) following the 3^rd^ seizure. Male and female C57BL6/J mice were not significantly different from each other (t(8)=1.11, p=0.30). (**E and F**) Mean oxygen profile before, during, and after MES in mixed sex *Slc1a3* (astrocyte reporter) *(E)* and *Pdgfrβ* (mural cell reporter) *(F)* mice (n=5, each). (**G**) Quantification (mean ± SEM) of the area below the severe hypoxic threshold (pO_2_<10mmHg) following the 3^rd^ seizure. *Slc1a3* and *Pdgfrβ* and combined male and female C57BL6/J mice were not significantly different from each other (F(2,17)=0.56, p=0.58). n.s.= not significant.

### Seizures cause sustained, COX-2 sensitive, arteriole constriction along with astrocytic and vascular smooth muscle Ca^2+^ recruitment

Severe cerebral hypoxic events are often the consequence of inadequate blood flow, which can occur via several mechanisms including sustained pathological vasoconstriction of contractile microvasculature, thrombotic or hemorrhagic stroke, a precipitous drop in perfusion pressure etc. Determining how local cerebral oxygenation and perfusion become dramatically decreased post epileptiform activity can identify new routes for intervention. We used two-photon fluorescence microscopy in awake mice and imaged single penetrating arterioles labelled with Rhod B-dextran in the barrel cortex to explore the mechanism of neocortical hypoperfusion/hypoxia. We first utlized *Slc1a3*-CreERT x RCL-GCaMP3 mice to observed astrocyte Ca^2+^ activity, along with microvascular responses, before, during and after a second MES-induced seizure (Figure 2 A,B). In response to 0.2 sec of MES, arterioles displayed a robust and prolonged vasoconstriction (ictal: −51.4 ± 8.5%, p<0.001; postictal at time 4800 sec: −37.7 ± 4.3%, p=0.007, n=5, Figure 2 C,D). The vasoconstriction lasted greater than 90 min in some trials, exhibiting a similar temporal profile to the deterioration and eventual recovery of oxygen levels. Even at peak constriction, the arteriole lumen could still be clearly visualized, and red blood cell movement detected, suggesting the vessel did not occlude. Examining astrocyte Ca^2+^ along with vascular lumen dynamics during and after seizure, we found that MES caused a rapid elevation in astrocyte endfoot Ca^2+^ during the initiation of vasoconstriction (361.7 ± 56.6%, p=0.007, n=4, Figure 2 C, E). The endfoot Ca^2+^ elevation decayed to a plateau (215.2 ± 61.5%, Figure 2 C,E) for ∼20 sec before returning to baseline values whereas vasoconstriction was sustained. Postictally at 4800 sec, endfoot Ca^2+^ was not different from baseline values (15.1 ± 13.5%, p=0.2, n=4, Figure 2 C,E), whereas lumen diameter was significantly smaller.

**Figure 2:**
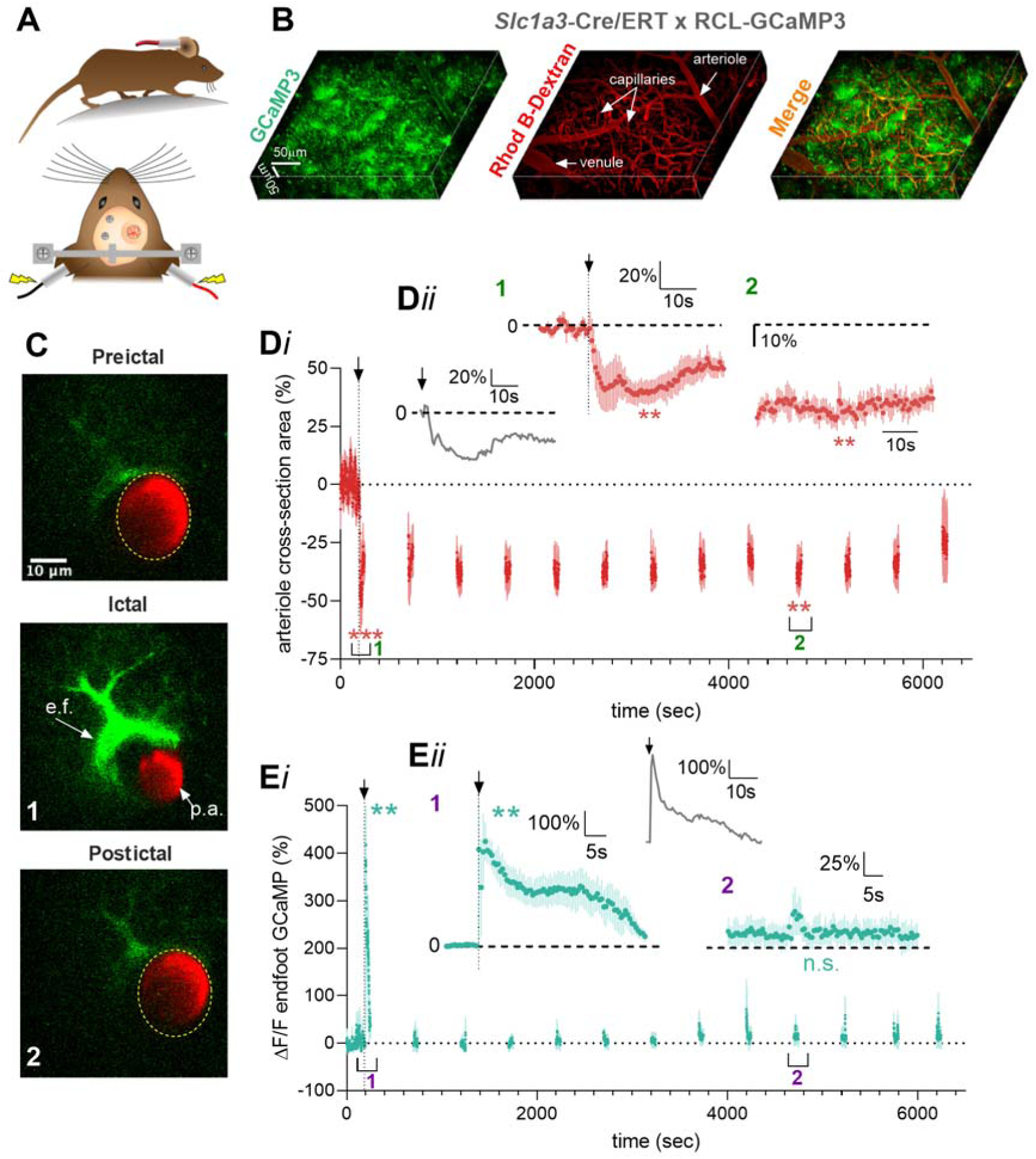
Seizure induced sustained arteriole constriction is associated with an initial transient rise in astrocytic endfoot Ca^2+^. (**A**) Schematics of awake mouse experimental setup. (**B**) 3D reconstruction of the superficial barrel cortex from a *Slc1a3*-Cre/ERT RCL-GCaMP3 mouse. Astrocytes expressing GCaMP3 are shown in green; the vasculature are loaded with Rhod B-dextran shown in red. (**C**) Cross section of a penetrating arteriole enwrapped by an endfoot (e.f.). Images show pre-ictal (top), severe vasoconstriction and a large astrocyte Ca^2+^ rise triggered by MES during the ictal period (middle), and the post-ictal period (bottom). (**D*i*)**. Summary time course of arteriolar diameter in *Slc1a3*-Cre/ERT RCL-GCaMP3 mice (n=5). To limit photobleaching/photodamage, measurements were taken every 300 sec for 60 sec. Arrow and vertical dotted line indicate MES (0.2sec). (**D*ii***) Temporal close-up of percent diameter changes during the ictal (t(4)=8.06, p<0.001) and postictal (t(4)=4.04, p=0.007) period. Inset: representative trace of diameter response to 0.2sec MES. (**E*i***) Summary time course of endfoot Ca^2+^ measurements in the same experiment as diameter measures (n=4). Vertical dotted line indicates MES (0.2s). (**E*ii***) Temporal close-up of endfoot Ca^2+^ response during the ictal (t(3)=6.36, p=0.007) and postictal (t(3)=1.63, p=0.20) period. Inset: representative trace of endfoot Ca^2+^ to 1sec MES. Data is presented as Mean ± S.E.M. **=p<0.01, ***=p<0.001. n.s.= not significant.

As we previously demonstrated a strong block of seizure-induced hypoxia by COX-2 antagonism, we tested whether ibuprofen (100 mg/kg) was effective at reducing sustained vasoconstriction. As ibuprofen is ineffective if administered shortly after seizure (Farrell 2016), indicating the necessity of ecosanoid production during epileptiform activity, we pretreated mice i.p. 30 minutes before imaging (Figure 3A). Notably, ibuprofen failed to block the initial rapid constriction that occurred from MES within the first ∼10 sec (ictal control: −45.8 ± 17.0% vs ictal ibuprofen: −34.7 ± 19.4, p=0.72), but COX-2 antagonism completely prevented the development of the prolonged vasoconstriction when compared to vehicle control (postictal vehicle at 5200sec: −38.7 ± 3.6% vs postictal ibuprofen at 5200 sec: −8.5 ± 5.3%, p=0.007, n=5, Figure 3 B,C). These data provide evidence that postictal hypoperfusion/hypoxia occurs via a COX-2 dependent enduring vasoconstriction in local microvasculature.

**Figure 3:**
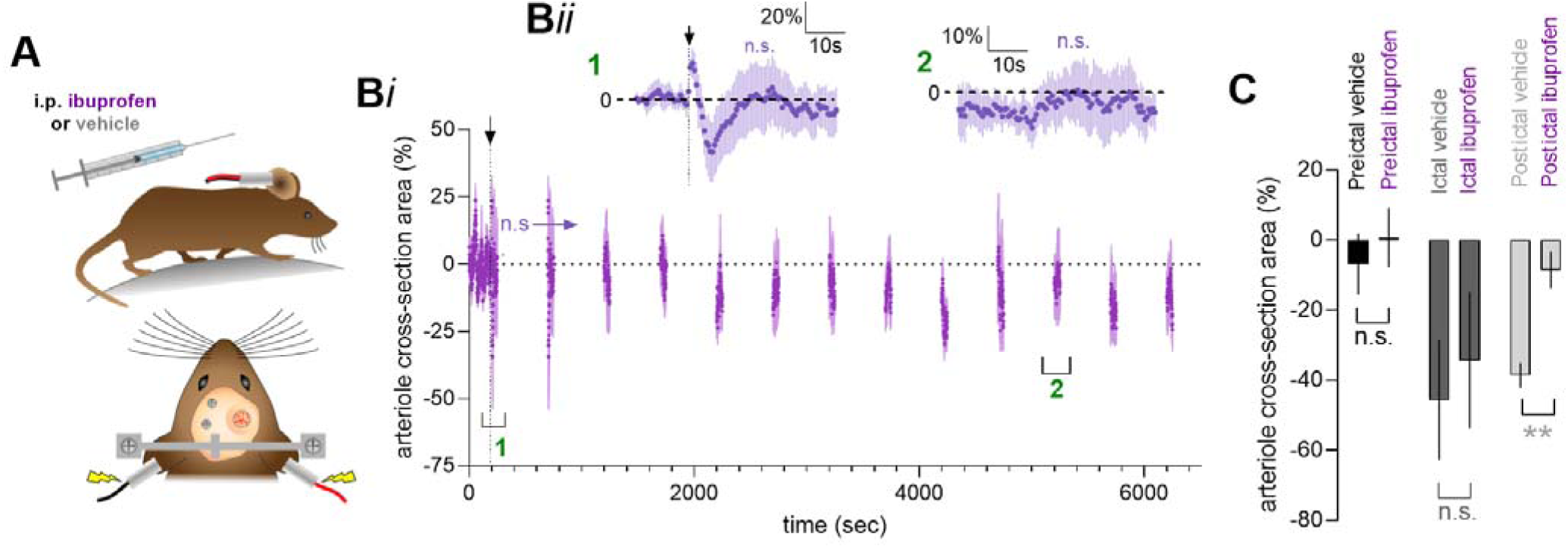
Postictal vasoconstriction is prevented by COX-2 antagonism. (**A**) Schematics of experimental setup with prior i.p. injection. (**B*i*).** Summary time course of arteriolar diameter in the presence of ibuprofen in response to MES (n=5). To limit photobleaching/photodamage, measurements were taken every 300 sec for 60 sec. Arrow and vertical dotted line indicates MES (0.2 sec). (**B*ii***) Temporal close-up of percent diameter changes during the ictal and postictal period (5200 sec vs baseline: t(4)=0.92, p=0.41). (**C**) Summary data comparing vehicle i.p. injection (grey) with ibuprofen (purple), at a pre-ictal (t(4)=1.31, p=0.25), ictal (t(4)=0.38, p=0.72) and post-ictal time point (5200 sec: t(4)=4.14, p=0.007). Data is presented as Mean ± S.E.M. **=p<0.01. n.s. = not significant.

Finally, we examined mural cell Ca^2+^ activity by using *Pdgfrβ*-Cre x RCL-GCaMP6s mice (n=5, Figure 4 A,B). By observing penetrating arterioles in response to MES, we found that VSMC Ca^2+^ exhibited a rapid increase in free Ca^2+^ (257.7 ± 108.9%, p=0.036, Figure 4 D,E) as the vessel constricted (−36.5 ± 7.6%, p=0.02, Figure 4 C,E), yet the Ca^2+^ signal maintained elevated above baseline values at various time points, even at 4500 sec (42.7 ± 15.4%, p=0.038) during the sustained vasoconstriction (−24.0 ± 7.6%, p=0.023). Furthermore, we examined the Spearman correlation coefficient between vascular smooth muscle Ca^2+^ and diameter using the averaged data from all 5 mice between baseline data compared to each ictal and postictal imaging epoch (each 30sec of data) and found significant negative relationships (Figure 4F*i*)(4500sec: r = −0.84, p<0.001, Figure 4 F*ii*). This suggested that VSMCs Ca^2+^ corresponded to the changes in arteriole diameter. Collectively, these data reveal the temporal dynamics of penetrating arteriole diameter, endfoot Ca^2+^ and VSMC Ca^2+^ during and up to 90min after seizure in awake mice.

**Figure 4:**
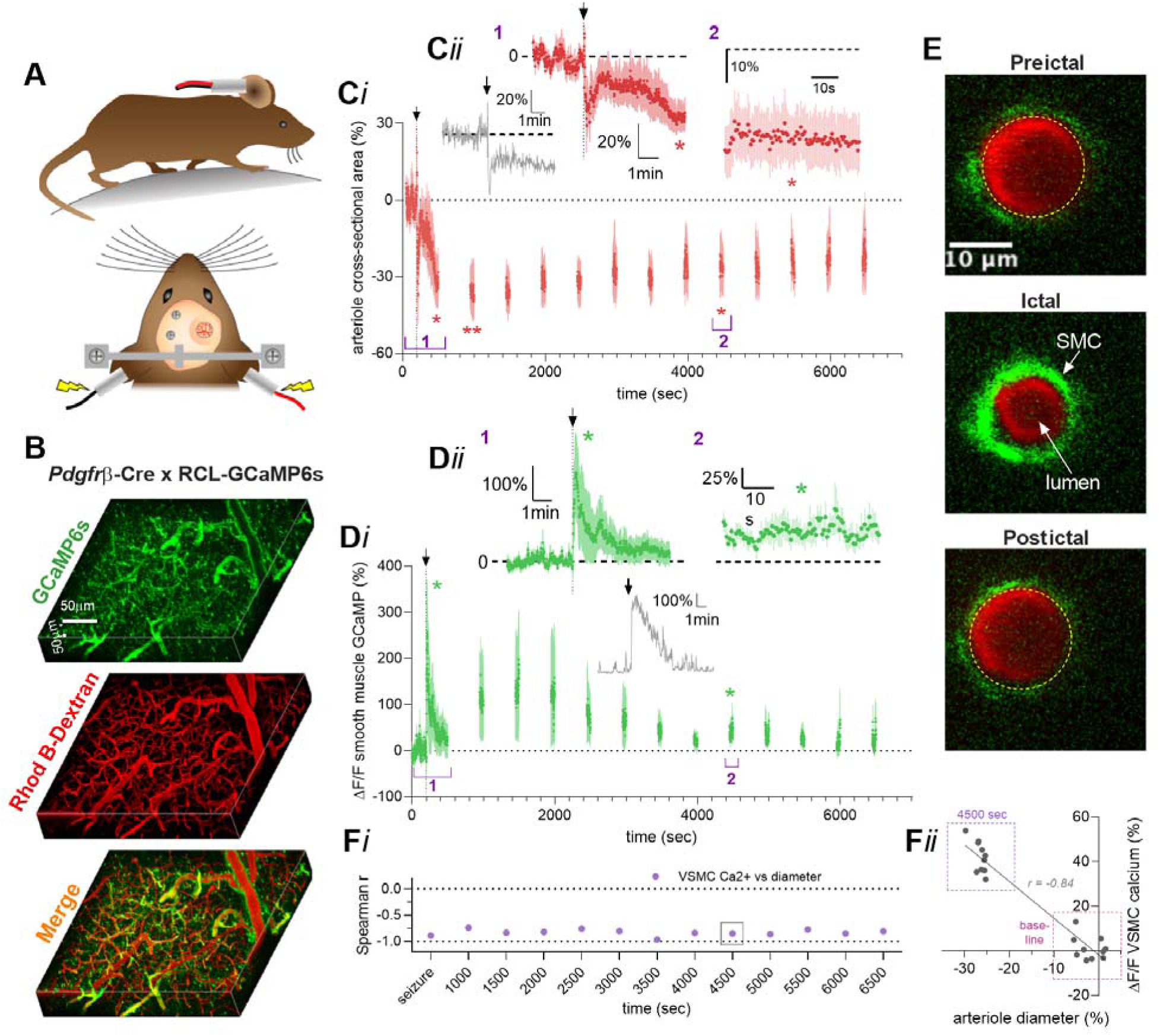
Seizure induced sustained arteriole constriction is associated with rapid and prolonged vascular smooth muscle cell Ca^2+^ elevation. (**A**) Schematics of awake mouse experimental setup. (**B**) 3D reconstruction of the superficial barrel cortex from a *Pdgfrβ*-Cre RCL-GCaMP6s mouse. VSMC expressing GCaMP6s are shown in green; the vasculature is loaded with Rhod B-dextran is shown in red. (**C*i***). Summary time course of arteriole diameter responses (n=5). Arrow and vertical dotted line indicate MES (0.2 sec). To limit photobleaching/photodamage, measurements were taken every 300 sec for 60 sec. (**C*ii***) Temporal close-up of percent diameter changes during the ictal (t(4)=3.74, p=0.02) and postictal (t(4)=3.58, p=0.023) period. Inset: representative trace of diameter in response to 0.2 s MES. (**D*i***) Summary time course of VSMC Ca^2+^ elevations in the same experiments as diameter measures. (**D*ii***) Temporal close-up of percent VSMC Ca^2+^ changes during the ictal (t(4)=2.42, p=0.036) and postictal (t(4)=3.03, p=0.038) period. Inset: representative trace of VSMC Ca^2+^ in response to 0.2 s MES. (**E**) Cross section of a penetrating arteriole (red) with VSMC expressing GCaMP6s (green). Images show baseline (top), the ictal period (middle) and the postictal period (bottom). (**F*i***) Summary of calculated Spearman r values between changes in arteriole diameter and VSMC Ca^2+^ over the postictal period. (**F*ii***) Correlation between VSMC Ca^2+^ and arteriole diameter during baseline (100sec before MES) and postictal period (4500sec). Each data point represents a 10sec bin and averaged across all 5 animals. Data is presented as Mean ± S.E.M. *=p<0.05, **=p<0.01.

## Discussion

Here we observed the calcium dynamics in astrocytes and vascular smooth muscle cells after brief, maximum electroconvulsive seizures along with corresponding changes in penetrating arteriole diameter. For the first time, we captured COX-2 sensitive, severe vasoconstriction in cerebral microvasculature in vivo that were sustained for over an hour after the ictal event. Astrocyte endfeet and vascular smooth muscle calcium signals were distinctly different in their temporal profile: endfoot calcium being restricted largely to the ictal period whereas smooth muscle calcium elevation was variable and lasted up to 75 minutes after MES. We also showed that postictal hypoxia generalizes equally to male and female mice and to the cre-lox knockin crosses used in our study.

We previously identified that COX-2 plays a central role in coordinating a cascade of events to induce severe hypoxia following electrographic seizures (Farrell et al., 2016). Here we show that COX-2 is necessary for pathological vasoconstrictions, which is most likely the primary cause for the hypoxia/hypoperfusion event. Activity-dependent induction of COX-2 in neurons plays an important role in normal neurovascular coupling by producing vasoactive prostanoids that act on blood vessel receptors to cause vasodilation (Lacroix et al., 2015; Lecrux et al., 2012; Lecrux et al., 2011). Under pathophysiological activation during electrographic seizures, we observed prolonged vasoconstriction rather than transient vasodilation. Though astrocytes have low levels of COX-2 under basal conditions, these cells up regulate COX-2 in models of epilepsy (Desjardins et al., 2003). Thus, the source of the constricting eicosanoids still needs to be delineated with cell-type selective COX-2 knockout experiments. For example, while seizure activity is expected to cause large increases in free calcium in principal neurons (Wenzel, Hamm, Peterka, & Yuste, 2017), we observed a large calcium signal also in endfeet which may have produced COX dependent vasoconstrictors. Prostanoids such as PGE2 that derive from COX activity can produce vasoconstriction if the appropriate EP receptors (EP1 and EP3) are present on vasculature (Dabertrand et al., 2013; Jadhav et al., 2004). Once released, our evidence indicates long-term changes to vascular smooth muscle calcium. Transient release of constrictor agents from parenchymal cells is all that is required as our previous data showed that COX-2 antagonism after seizure was ineffective, whereas postictal administration of an L-type antagonist was effective at blocking the sustained drop in tissue oxygen (Farrell et al., 2016). How vascular calcium elevation becomes sustained is still unclear, but a mechanism that acts to facilitate L-type Ca^2+^ channel opening and/or decrease plasmalemma K^+^ channel opening, which help set resting membrane potential thereby affect L-type recruitment, are possibilities. Our observations will help aid future experiments to investigate these ideas.

Previously we had used acute hippocampal slices and observed the diameter of local arterioles (Farrell et al., 2016). The conditions in slice differ substantially from *in vivo* conditions, especially regarding the lack of blood pressure and flow, and this could potentially misrepresent the arteriolar results. For example, slices neither capture vascular temporal dynamics accurately, nor are the magnitude of the arteriole diameter measurements compared to *in vivo* (Filosa, 2010). However, the degree of constriction observed in slice provided approximate estimates to the changes in blood perfusion that were observed with laser Doppler flowmetry. Furthermore, the vasoconstriction following seizure stimulation in slice was sensitive to acetaminophen and nifedipine, which supports the notion that this constriction follows the same set of mechanisms that occur *in vivo*. Nevertheless, our *in vivo* observations of arterioles were essential to either support or refute our current model.

Postictal behavioral impairments last much longer than the seizures themselves and negatively impact quality of life (Josephson et al., 2016). The mechanism underlying these disruptions were previously unknown and no treatment options existed. Our discovery that brief and mild seizures lead to an extended vasoconstrictive event is critically important because it establishes that seizures could injure the brain through *postictal hypoperfusion/hypoxia*, and not necessarily by the seizure itself. Severe hypoxia may be an important component of seizure-induced brain damage (Jackson, Chambers, & Berkovic, 1999) and since postictal hypoperfusion/hypoxia is COX-2 dependent, this hypothesis can be assessed. Though there is some controversy (Rojas et al., 2014), it is generally accepted that COX-2 inhibition is neuroprotective from seizures. Indeed, with genetic or pharmacological COX-2 inhibition, seizure-induced brain damage can be dramatically reduced (Hewett, Silakova, & Hewett, 2006; Kunz & Oliw, 2001; Serrano et al., 2011). Though at the time of these studies researchers were unaware of postictal hypoperfusion/hypoxia and the requirement of COX-2 in this response, they provided support for the hypothesis that postictal hypoperfusion/hypoxia injures the brain (Farrell et al., 2017). Given the central role of brain injury in epileptogenesis (Sloviter & Bumanglag, 2013), preventing injury from vasoconstriction-induced hypoperfusion/hypoxia may be a novel preventative treatment strategy in epilepsy.

## Materials and methods

### Mice

Mice were handled and maintained according to the Canadian Council for Animal Care guidelines. These procedures were approved by the Life and Environmental Sciences Animal Care and Health Sciences Animal Care Committees at the University of Calgary (AC15-0133) (AC16-0272). All studies were either performed on young adult male and female C57BL6/J, *Slc1a3*-Cre/ERT (Jax:012586) × RCL-GCaMP3 (Jax: 014538, Ai38) and *Pdgfrβ*-Cre (courtesy of Volkard Lindner and Andy Shih) × RCL-GCaMP6s (Jax: 024106, Ai96) mice weighing between 21-30g. Mice were housed individually in clear plastic cages and were maintained on a 12hr: 12hr light/dark cycle lights on at 07:00 hours, in separate colony rooms under specified pathogen free conditions. Food and water were available *ad libitum*. All experimental procedures occurred during the light phase.

### Eliciting and Recording Seizures and Oxygen Detection

Electrodes were constructed from Teflon-coated, stainless steel wire, 178 µm in diameter (A-M Systems). Wire ends were stripped of Teflon and connected to gold-plated male amphenol pins. Mice were anaesthetized with a 4% isoflurane and maintained between 1% and 2%. Lidocaine (2%) was administered subcutaneously at the incision site. One bipolar electrode was chronically implanted under stereotaxic control in barrel cortex at −1.75 AP, +3 ML, −1.5 DV with an oxygen-sensing probe positioned nearby at −1.65, AP, +3.5 ML, −1.5 DV relative to bregma. The implants were adhered and anchored to the skull using dental cement and a ground electrode. Subsequent experimental procedures commenced no earlier than 5 days following surgery.

Oxygen recordings were obtained using an implantable fiber-optic oxygen-sensing device. 525nm light pulses induce fluorescence (measured at 650nm) at the platinum tip that is quenched by oxygen within a local area (∼500 µm^3^) and uses the fluorescence decay time to derive pO_2_ (Ortiz-Prado et al., 2010). The technology (Oxylite, Oxford Optronics) does not consume oxygen while measuring absolute pO_2_ values. The manufacturer individually calibrates each biologically inert probe, called an optrode. The implant is inserted under isoflurane anesthesia, and every effort was made to minimize suffering. We allow at least 7 days between implantation and initiation of measurements to ensure that the effects of acute trauma were minimized. pO_2_ measurements at 1Hz can then be made at any time by connecting the implant to the Oxylite using an extension fiber optic lead. The probe provides accurate and continuous measurements of local pO_2_ levels in brain tissue in awake, freely moving animals over several weeks and is exquisitely sensitive to oxygen perturbations in relation to epileptiform activity (Farrell, Greba, Snutch, Howland, & Teskey, 2018).

On test days, mice were connected to the EEG, oxygen-sensing system and saline soaked ear clips and allowed 5 minutes to adjust before any measurements were taken. A seizure was elicited after 100 seconds of baseline recording using a suprathreshold MES stimulus, which was delivered through the ear clips via a GSC 700 shock generator (model E1100DA) (Grason-Sradler). Oxygen levels were recorded before and after the delivery of a 0.2 second train of 60 Hz biphasic sine-wave pulses (Young et al., 2006). Seizure duration was recorded by observing seizure behavior (Racine, 1972). Once the EEG returned to baseline following a seizure, the electrodes were disconnected, but the fiber-optic cable for oxygen-sensing was left attached. The fiber-optic cable was disconnected when pO_2_ levels returned to baseline.

### Awake In Vivo Two-Photon Preparation

All surgeries followed the procedures as detailed as previously described (Tran & Gordon, 2015a). Briefly, one week before the imaging session, a custom head bar was surgically installed without performing a craniectomy and the animal was then returned to a new home cage to recover for two days (single housing). The mouse was then trained on a passive air-supported Styrofoam ball treadmill with its head restrained. The training was done on two consecutive days for 30min session each. The animals were habituated to the seizure setup without inducing the actual seizure (i.e. the ears were clipped with the banana clips). The animals were allowed to return to their home cage after each training session. On the second day the mouse received MES-induced seizure during ball training. On the imaging day, a cranial window was created as previously described over the primary somatosensory cortex with both bone and dura removed.

### Vessel Indicators

Rhodamine B isothiocyanate (RhodB)-dextran (MW 70,000, Sigma) was tail vein injected (100-200mL of 2.3% (w/v) solution in saline) to visualize the blood plasma. Before imaging the mouse recovered on the treadmill with its head immobilized for 30 minutes.

### Two-Photon Fluorescence Microscopy

Fluorescence images were obtained using a custom built *in vivo* two-photon microscope (Rosenegger, Tran, LeDue, Zhou, & Gordon, 2014) fed by a tunable Ti:sapphire laser (Coherent Chameleon, Ultra II, ∼4Wavg power, 670-1080 nm, ∼80 MHz, 140 fs pulse width), equipped with GaAsP PMTs (Hamamatsu, Japan) and controlled by open-source ScanImage software (https://wiki.janelia.org/). We used a Nikon 16X, 0.8NA, 3mm WD objective lens or a Zeiss 40X, 1.0NA, 2.5mm WD objective lens. GCaMP3 was excited at 920 nm. Green fluorescence signals were filtered using a 525/50nm BP, and orange/red light was filtered using a 605/70nm BP (Chroma Technology, Bellows Falls VT USA). Bidirectional xy raster scanning was used at a frame rate of 0.98 Hz.

### Behavior Capture

A near infrared LED (780 nm) and camera were used to capture simple behaviors such as resting, running, whisking and tracking animal responding to MES concurrently with two-photon fluorescence imaging for all experimental trials at a frame rate of 14 Hz.

### Seizure Induction for Two-photon Imaging

On imaging days, mice were connected to the ear clips and allowed 5 minutes for the animals to adjust before imaging. A seizure was elicited after 5 minutes of baseline imaging using a suprathreshold MES stimulus that was delivered through the ear clips via a GSC 700 shock generator (model E1100DA) (Grason-Sradler). Continuous imaging was conducted for 15 minutes and 1 minute for every 5 minutes afterward.

### Replicates

For technical replicates (a test performed on the same sample multiple times), whether recording oxygen and EEG or performing two-photon imaging, each animal received three MES total. For biological replicates (a test performed on biologically distinct samples representing an identical time point or treatment dose), we performed oxygen and EEG measures on four distinct groups of mice: male C57Bl/6J, female C57Bl/6J, Astrocyte Ca2+ reporter mice and VSMC Ca2+ reporter mice, and observed the same postictal hypoxia in all groups. We also performed the same two-photon imaging procedure on two groups: Astrocyte Ca2+ reporter mice and VSMC Ca2+ reporter mice and observed the same postictal vasoconstriction in both groups. Mice that did not exhibit behavioral seizures to stimulation were excluded from the study. Furthermore, we did not excluded outliers. Power calculations were not performed to determine sample size. Mice were obtained from our in-house breeding colonies and each served as its own control.

### Statistics

All statistical analyses were performed using Prism version 5.01 (GraphPad). Statistical ‘n’ represented the measurements from a given mouse. One or two tailed paired, parametric t-tests were used for experiments with only two groups. One tailed t-tests were used for ibuprofen vs vehicle on arteriole diameter measures, as well as VSMC Ca^2+^ changes vs baseline, as our previous published data allowed use to predict the direction of the effect. ANOVAs were used for experiments with more than 2 groups and a follow-up Tukey test to identify in which group(s) the significant differences occur. Repeated measures statistics were used for all within subject experiments. Spearman’s correlation coefficient ‘r’ was used to look at the relationship between VSMC Ca^2+^ and arteriole diameter.

## Acknowledgments

This work was supported by the Canada Institutes of Health Research. G.R.G. was also supported by Canada Research Chairs. A.G. was supported by SUDEP Aware. We thank Dr. Volkhard Lindner at the University of Tübingen, care of Dr. Andy Shih at the University of Seattle for supplying us with PDGFRβ-Cre mice. We thank the developers and distributers of ScanImage open-source control and acquisition software for two-photon laser scanning microscopy.

## Author Contributions

Conceptualization, G.C.T., G.R.G; Investigation, C.H.T, A.G.G.; Writing – Original Draft, all authors; Writing – Review and Editing, all authors; Supervision, G.R.G., and G.C.T.; Funding: G.R.G. (148471) and G.C.T. (MOP-130495).

## References

Bondy, S. C., Mitchell, C. L., Rahmaan, S., & Mason, G. (1987). Regional variation in the response of cerebral ornithine decarboxylase to electroconvulsive shock. Neurochemical Pathology, 7(2), 129–141.

Cuttler, A. S., LeClair, R. J., Stohn, J. P., Wang, Q., Sorenson, C. M., Liaw, L., & Lindner, V. (2011). Characterization of Pdgfrb-Cre transgenic mice reveals reduction of ROSA26 reporter activity in remodeling arteries. Genesis (New York, N.Y.: 2000), 49(8), 673–680. https://doi.org/10.1002/dvg.20769

Dabertrand, F., Hannah, R. M., Pearson, J. M., Hill-Eubanks, D. C., Brayden, J. E., & Nelson, M. T. (2013). Prostaglandin E2, a postulated astrocyte-derived neurovascular coupling agent, constricts rather than dilates parenchymal arterioles. Journal of Cerebral Blood Flow and Metabolism: Official Journal of the International Society of Cerebral Blood Flow and Metabolism, 33(4), 479–482. https://doi.org/10.1038/jcbfm.2013.9

Desjardins, P., Sauvageau, A., Bouthillier, A., Navarro, D., Hazell, A. S., Rose, C., & Butterworth, R. F. (2003). Induction of astrocytic cyclooxygenase-2 in epileptic patients with hippocampal sclerosis. Neurochemistry International, 42(4), 299–303.

Erecinska, M., & Silver, I. A. (2001). Tissue oxygen tension and brain sensitivity to hypoxia. Respiration Physiology, 128(3), 263–276.

Farrell, J. S., Colangeli, R., Wolff, M. D., Wall, A. K., Phillips, T. J., George, A., … Teskey, G. C. (2017). Postictal hypoperfusion/hypoxia provides the foundation for a unified theory of seizure-induced brain abnormalities and behavioral dysfunction. Epilepsia, 58(9), 1493–1501. https://doi.org/10.1111/epi.13827

Farrell, J. S., Gaxiola-Valdez, I., Wolff, M. D., David, L. S., Dika, H. I., Geeraert, B. L., … Teskey, G. C. (2016). Postictal behavioural impairments are due to a severe prolonged hypoperfusion/hypoxia event that is COX-2 dependent. ELife, 5. https://doi.org/10.7554/eLife.19352

Farrell, J. S., Greba, Q., Snutch, T. P., Howland, J. G., & Teskey, G. C. (2018). Fast oxygen dynamics as a potential biomarker for epilepsy. Scientific Reports, 8(1), 17935. https://doi.org/10.1038/s41598-018-36287-2

Filosa, J. A. (2010). Vascular tone and neurovascular coupling: considerations toward an improved in vitro model. Frontiers in Neuroenergetics, 2. https://doi.org/10.3389/fnene.2010.00016

Gaxiola-Valdez, I., Singh, S., Perera, T., Sandy, S., Li, E., & Federico, P. (2017). Seizure onset zone localization using postictal hypoperfusion detected by arterial spin labelling MRI. Brain: A Journal of Neurology, 140(11), 2895–2911. https://doi.org/10.1093/brain/awx241

Gordon, G. R. J., Choi, H. B., Rungta, R. L., Ellis-Davies, G. C. R., & MacVicar, B. A. (2008). Brain metabolism dictates the polarity of astrocyte control over arterioles. Nature, 456(7223), 745–749. https://doi.org/10.1038/nature07525

Grant, R. I., Hartmann, D. A., Underly, R. G., Berthiaume, A.-A., Bhat, N. R., & Shih, A. Y. (2019). Organizational hierarchy and structural diversity of microvascular pericytes in adult mouse cortex. Journal of Cerebral Blood Flow and Metabolism: Official Journal of the International Society of Cerebral Blood Flow and Metabolism, 39(3), 411–425. https://doi.org/10.1177/0271678X17732229

Hewett, S. J., Silakova, J. M., & Hewett, J. A. (2006). Oral treatment with rofecoxib reduces hippocampal excitotoxic neurodegeneration. The Journal of Pharmacology and Experimental Therapeutics, 319(3), 1219–1224. https://doi.org/10.1124/jpet.106.109876

Hill, R. A., Tong, L., Yuan, P., Murikinati, S., Gupta, S., & Grutzendler, J. (2015). Regional Blood Flow in the Normal and Ischemic Brain Is Controlled by Arteriolar Smooth Muscle Cell Contractility and Not by Capillary Pericytes. Neuron, 87(1), 95–110. https://doi.org/10.1016/j.neuron.2015.06.001

Höckel, M., & Vaupel, P. (2001). Biological consequences of tumor hypoxia. Seminars in Oncology, 28(2 Suppl 8), 36–41.

Howarth, C. (2014). The contribution of astrocytes to the regulation of cerebral blood flow. Frontiers in Neuroscience, 8, 103. https://doi.org/10.3389/fnins.2014.00103

Jackson, G. D., Chambers, B. R., & Berkovic, S. F. (1999). Hippocampal sclerosis: development in adult life. Developmental Neuroscience, 21(3–5), 207–214. https://doi.org/10.1159/000017400

Jadhav, V., Jabre, A., Lin, S.-Z., & Lee, T. J.-F. (2004). EP1- and EP3-receptors mediate prostaglandin E2-induced constriction of porcine large cerebral arteries. Journal of Cerebral Blood Flow and Metabolism: Official Journal of the International Society of Cerebral Blood Flow and Metabolism, 24(12), 1305–1316. https://doi.org/10.1097/01.WCB.0000139446.61789.14

Jiang, B. H., Semenza, G. L., Bauer, C., & Marti, H. H. (1996). Hypoxia-inducible factor 1 levels vary exponentially over a physiologically relevant range of O2 tension. The American Journal of Physiology, 271(4 Pt 1), C1172–1180. https://doi.org/10.1152/ajpcell.1996.271.4.C1172

Josephson, C. B., Engbers, J. D. T., Sajobi, T. T., Jette, N., Agha-Khani, Y., Federico, P., … Wiebe, S. (2016). An investigation into the psychosocial effects of the postictal state. Neurology, 86(8), 723–730. https://doi.org/10.1212/WNL.0000000000002398

Kunz, T., & Oliw, E. H. (2001). The selective cyclooxygenase-2 inhibitor rofecoxib reduces kainate-induced cell death in the rat hippocampus. The European Journal of Neuroscience, 13(3), 569–575.

Lacroix, A., Toussay, X., Anenberg, E., Lecrux, C., Ferreirόs, N., Karagiannis, A., … Cauli, B. (2015). COX-2-Derived Prostaglandin E2 Produced by Pyramidal Neurons Contributes to Neurovascular Coupling in the Rodent Cerebral Cortex. The Journal of Neuroscience: The Official Journal of the Society for Neuroscience, 35(34), 11791–11810. https://doi.org/10.1523/JNEUROSCI.0651-15.2015

Lecrux, C., Kocharyan, A., Sandoe, C. H., Tong, X.-K., & Hamel, E. (2012). Pyramidal cells and cytochrome P450 epoxygenase products in the neurovascular coupling response to basal forebrain cholinergic input. Journal of Cerebral Blood Flow and Metabolism: Official Journal of the International Society of Cerebral Blood Flow and Metabolism, 32(5), 896– 906. https://doi.org/10.1038/jcbfm.2012.4

Lecrux, C., Toussay, X., Kocharyan, A., Fernandes, P., Neupane, S., Lévesque, M., … Hamel, E. (2011). Pyramidal neurons are “neurogenic hubs” in the neurovascular coupling response to whisker stimulation. The Journal of Neuroscience: The Official Journal of the Society for Neuroscience, 31(27), 9836–9847. https://doi.org/10.1523/JNEUROSCI.4943-10.2011

Li, E., d’Esterre, C.D., Gaxiola-Valdez, I., Lee, T-Y., Menon, B., Peedicail, J.S.. … P. Federico. (2019) CT perfusion measurement of postictal hypoperfusion: localization of the seizure onset zone and patterns of spread. Neuroradiology accepted.

Maloney-Wilensky, E., Gracias, V., Itkin, A., Hoffman, K., Bloom, S., Yang, W., … LeRoux, P. D. (2009). Brain tissue oxygen and outcome after severe traumatic brain injury: a systematic review. Critical Care Medicine, 37(6), 2057–2063. https://doi.org/10.1097/CCM.0b013e3181a009f8

Mehina, E. M. F., Murphy-Royal, C., & Gordon, G. R. (2017). Steady-State Free Ca2+ in Astrocytes Is Decreased by Experience and Impacts Arteriole Tone. The Journal of Neuroscience: The Official Journal of the Society for Neuroscience, 37(34), 8150–8165. https://doi.org/10.1523/JNEUROSCI.0239-17.2017

Mishra, A. (2017). Binaural blood flow control by astrocytes: listening to synapses and the vasculature. The Journal of Physiology, 595(6), 1885–1902. https://doi.org/10.1113/JP270979

Mulligan, S. J., & MacVicar, B. A. (2004). Calcium transients in astrocyte endfeet cause cerebrovascular constrictions. Nature, 431(7005), 195–199. https://doi.org/10.1038/nature02827

Neppl, R. L., Lubomirov, L. T., Momotani, K., Pfitzer, G., Eto, M., & Somlyo, A. V. (2009). Thromboxane A2-induced bi-directional regulation of cerebral arterial tone. The Journal of Biological Chemistry, 284(10), 6348–6360. https://doi.org/10.1074/jbc.M807040200

Nishimura, N., Schaffer, C. B., Friedman, B., Lyden, P. D., & Kleinfeld, D. (2007). Penetrating arterioles are a bottleneck in the perfusion of neocortex. Proceedings of the National Academy of Sciences of the United States of America, 104(1), 365–370. https://doi.org/10.1073/pnas.0609551104

Ortiz-Prado, E., Natah, S., Srinivasan, S., & Dunn, J. F. (2010). A method for measuring brain partial pressure of oxygen in unanesthetized unrestrained subjects: the effect of acute and chronic hypoxia on brain tissue PO(2). Journal of Neuroscience Methods, 193(2), 217–225. https://doi.org/10.1016/j.jneumeth.2010.08.019

Racine, R. J. (1972). Modification of seizure activity by electrical stimulation. II. Motor seizure. Electroencephalography and Clinical Neurophysiology, 32(3), 281–294.

Rojas, A., Jiang, J., Ganesh, T., Yang, M.-S., Lelutiu, N., Gueorguieva, P., & Dingledine, R. (2014). Cyclooxygenase-2 in epilepsy. Epilepsia, 55(1), 17–25. https://doi.org/10.1111/epi.12461

Rosenegger, D. G., Tran, C. H. T., LeDue, J., Zhou, N., & Gordon, G. R. (2014). A high performance, cost-effective, open-source microscope for scanning two-photon microscopy that is modular and readily adaptable. PloS One, 9(10), e110475. https://doi.org/10.1371/journal.pone.0110475

Sakadžic, S., Yaseen, M. A., Jaswal, R., Roussakis, E., Dale, A. M., Buxton, R. B., … Devor, A. (2016). Two-photon microscopy measurement of cerebral metabolic rate of oxygen using periarteriolar oxygen concentration gradients. Neurophotonics, 3(4), 045005. https://doi.org/10.1117/1.NPh.3.4.045005

Sato, M., Tani, E., Fujikawa, H., & Kaibuchi, K. (2000). Involvement of Rho-kinase-mediated phosphorylation of myosin light chain in enhancement of cerebral vasospasm. Circulation Research, 87(3), 195–200.

Serrano, G. E., Lelutiu, N., Rojas, A., Cochi, S., Shaw, R., Makinson, C. D., … Dingledine, R. (2011). Ablation of cyclooxygenase-2 in forebrain neurons is neuroprotective and dampens brain inflammation after status epilepticus. The Journal of Neuroscience: The Official Journal of the Society for Neuroscience, 31(42), 14850–14860. https://doi.org/10.1523/JNEUROSCI.3922-11.2011

Slezak, M., Göritz, C., Niemiec, A., Frisén, J., Chambon, P., Metzger, D., & Pfrieger, F. W. (2007). Transgenic mice for conditional gene manipulation in astroglial cells. Glia, 55(15), 1565–1576. https://doi.org/10.1002/glia.20570

Sloviter, R. S., & Bumanglag, A. V. (2013). Defining “epileptogenesis” and identifying “antiepileptogenic targets” in animal models of acquired temporal lobe epilepsy is not as simple as it might seem. Neuropharmacology, 69, 3–15. https://doi.org/10.1016/j.neuropharm.2012.01.022

Srienc, A. I., Kornfield, T. E., Mishra, A., Burian, M. A., & Newman, E. A. (2012). Assessment of glial function in the in vivo retina. Methods in Molecular Biology (Clifton, N.J.), 814, 499–514. https://doi.org/10.1007/978-1-61779-452-0_33

Straub, S. V., Bonev, A. D., Wilkerson, M. K., & Nelson, M. T. (2006). Dynamic inositol trisphosphate-mediated calcium signals within astrocytic endfeet underlie vasodilation of cerebral arterioles. The Journal of General Physiology, 128(6), 659–669. https://doi.org/10.1085/jgp.200609650

Toda, N. (1982). Mechanism of action of carbocyclic thromboxane A2 and its interaction with prostaglandin I2 and verapamil in isolated arteries. Circulation Research, 51(6), 675–682.

Tran, C. H. T., & Gordon, G. R. (2015a). Acute two-photon imaging of the neurovascular unit in the cortex of active mice. Frontiers in Cellular Neuroscience, 9, 11. https://doi.org/10.3389/fncel.2015.00011

Tran, C. H. T., & Gordon, G. R. (2015b). Astrocyte and microvascular imaging in awake animals using two-photon microscopy. Microcirculation (New York, N.Y.: 1994), 22(3), 219–227. https://doi.org/10.1111/micc.12188

Tran, C. H. T., Peringod, G., & Gordon, G. R. (2018). Astrocytes Integrate Behavioral State and Vascular Signals during Functional Hyperemia. Neuron, 100(5), 1133-1148.e3. https://doi.org/10.1016/j.neuron.2018.09.045

van den Brink, W. A., van Santbrink, H., Steyerberg, E. W., Avezaat, C. J., Suazo, J. A., Hogesteeger, C., … Maas, A. I. (2000). Brain oxygen tension in severe head injury. Neurosurgery, 46(4), 868–876; discussion 876-878.

Wenzel, M., Hamm, J. P., Peterka, D. S., & Yuste, R. (2017). Reliable and Elastic Propagation of Cortical Seizures In Vivo. Cell Reports, 19(13), 2681–2693. https://doi.org/10.1016/j.celrep.2017.05.090

Young, N. A., Teskey, G. C., Henry, L. C., & Edwards, H. E. (2006). Exogenous antenatal glucocorticoid treatment reduces susceptibility for hippocampal kindled and maximal electroconvulsive seizures in infant rats. Experimental Neurology, 198(2), 303–312. https://doi.org/10.1016/j.expneurol.2005.11.013

Zhang, C., Tabatabaei, M., Bélanger, S., Girouard, H., Moeini, M., Lu, X., & Lesage, F. (2019). Astrocytic endfoot Ca2+; correlates with parenchymal vessel responses during 4-AP induced epilepsy: An in vivo two-photon lifetime microscopy study. Journal of Cerebral Blood Flow and Metabolism: Official Journal of the International Society of Cerebral Blood Flow and Metabolism, 39(2), 260–271. https://doi.org/10.1177/0271678X17725417

